# Spreading depolarization causes reperfusion failure after cerebral ischemia

**DOI:** 10.1101/2022.09.05.506665

**Authors:** Anna Törteli, Réka Tóth, Sarah D. Berger, Sarah Samardzic, Ferenc Bari, Ákos Menyhárt, Eszter Farkas

## Abstract

**Background:** Despite successful recanalization to treat acute ischemic stroke, reperfusion failure associated with poor functional outcomes develops in half of the patients. The cause of reperfusion failure remains the subject of intensive research. Here, we explore the possibility that spreading depolarization (SD), a potent ischemic injury mechanism is a significant contributor and reliable predictor of reperfusion failure.

**Methods:** Young adult male and female C57BL/6 mice (n=69) were anesthetized with isoflurane (0.6-0.9%) and prepared for transcranial optical imaging. After 10 min of baseline, incomplete global forebrain ischemia was induced by transient (45 min) bilateral common carotid artery (CCA) occlusion, followed by 75 min reperfusion. SD and cerebral blood flow (CBF) changes were visualized with intrinsic optical signal imaging and laser speckle contrast imaging. To block SD, the irreversible NMDA receptor antagonist MK801 was applied (0.3 mg/kg, i.p., n=29). Neurological deficit was evaluated at baseline and post-ischemia with a composite Garcia Neuroscore scale. Collaterals of the circle of Willis were examined after loading the vasculature with carbon black ink. Ischemic neuronal injury was evaluated in hematoxylin-eosin-stained brain sections.

**Results:** SD emerged after ischemia onset in one or both hemispheres under a perfusion threshold (CBF drop to 21.1±4.6 vs. 33.6±4.4 %, SD vs. no SD). The failure of later reperfusion (44.4±12.5 %) was invariably linked to previous ischemic SD. In contrast, reperfusion was adequate (98.9±7.4 %) in hemispheres devoid of SD during ischemia. CBF reduced below the perfusion threshold of SD, when the P1 segment was absent in the circle of Willis. SD occurrence and the linked reperfusion failure were associated with poor neurologic function, and greater neuronal necrosis. The inhibition of SD with MK801 significantly improved reperfusion.

**Conclusions:** SD occurrence during ischemia impairs later reperfusion, prognosticating poor functional outcomes. The increased likelihood of SD occurrence is predicted by inadequate collaterals.

## Introduction

Recanalization is the only therapeutic choice to treat acute ischemic stroke and is one of the strongest predictors of favorable outcome and reduced mortality rate^1^. However, in almost half of the patients, recanalization is futile, and there is no neurological improvement despite optimal angiographic results^2,3^. Futile recanalization or reperfusion failure is associated with older age and poor collaterals^1,2^. Despite frequent diagnosis, the mechanistic background of reperfusion failure has remained unresolved. The pathomechanisms suspected to account for reperfusion failure include microvascular no-reflow due to embolization, neutrophil obstruction or pericyte constriction, next to large vessel constriction^4^. In our present work, we explore the possibility that ischemic spreading depolarization (SD), a potent pathophysiologic phenomenon is a reliable predictor of reperfusion failure.

SD develops in a recurrent pattern in the ischemic cortex of malignant hemispheric stroke patients as recorded during decompressive hemicraniectomy^5^ or over the first week after the surgical procedure^6-8^. SD occurs first within minutes after the onset of vascular occlusion^9^. SD represents the disruption of the neuronal resting membrane potential in a critical volume of gray matter, which propagates as a wave over the nervous tissue^10^. The pathophysiological relevance of SD is thought to lie in the formation of cytotoxic edema, the accumulation of glutamate, and the building up of a massive acid load in the wake of the wavefront, which initiate intracellular cell death cascades^11-13^. In the acutely injured human cortex, specific SD patterns have been associated with lesion maturation and the development of delayed infarctions^14,15^. These reports and accumulating experimental evidence collectively testify that SDs spontaneously evolve during acute brain injury and facilitate infarct growth^16^. However, any putative relationship between SD and reperfusion failure has remained unattended.

Here we set out to demonstrate in the mouse cortex that SD that occurs during an ischemic episode has a profound impact on the success of subsequent reperfusion.

## Materials and methods

### Animals

The experimental procedures were approved by the National Food Chain Safety and Animal Health Directorate of Csongrád-Csanád County, Hungary. The procedures were performed according to the guidelines of the Scientific Committee of Animal Experimentation of the Hungarian Academy of Sciences [updated Law and Regulations on Animal Protection: 40/2013. (II. 14.) Gov. of Hungary], following the EU Directive 2010/63/EU on the protection of animals used for scientific purposes, and reported in compliance with the ARRIVE guidelines. The mice - obtained from the Charles River colony at the Biological Research Centre, Centre of Excellence of the European Union, Szeged, Hungary – were group-housed and kept under normal 12-h light/dark cycle, with *ad libitum* access to food and tap water.

### Surgical procedures

Young adult (10-12 weeks old) male (n= 35) and female (n= 34) C57BL/6 mice (m=25-31 g) were anesthetized with isoflurane (0.6-0.9% in N_2_O:O_2_) and allowed to breath spontaneously through a nose cone. Body temperature was maintained at 37 °C using a thermostat-controlled heating pad (CODA monitor, Kent Scientific Corp., Torrington, CT, USA). Topical Lidocaine (1%) was administered at incisions. The CCAs were separated, and silk threads were lopped around the arteries for later occlusion. Mice were then fixed in the prone position in a stereotactic frame. A midline incision was made above the sagittal suture, and the skin was gently pulled aside. The entire skull surface was covered with UV light adhesive (UV683 Light Curing Adhesive, Permabond Ltd., Wessex, UK) to increase transparency and to avoid drying out.

### Intrinsic optical signal (IOS) imaging and laser speckle contrast analysis (LASCA) through the intact skull

SD propagation was tracked by IOS imaging, while cerebral blood flow (CBF) changes were monitored with LASCA^17^. The entire cortical surface (both hemispheres) was illuminated in a stroboscopic mode with a light emitting diode (LED) (530 nm peak wavelength; SLS-0304-A, Mightex Systems, Pleasanton, CA, USA) and a laser diode (HL6545MG, Thorlabs Inc., New Jersey, USA; 120 mW; 660 nm emission wavelength) driven by a power supply (LDTC0520, Wavelength Electronics, Inc., Bozeman, USA) set to deliver a 160-mA current. Both image sequences were captured with a monochrome CCD camera (resolution: 1024 × 1024 pixel, Pantera 1M30, DALSA, Gröbenzell, Germany) attached to a stereomicroscope (MZ12.5, Leica Microsystems, Wetzlar, Germany). LED (100 ms/s) and laser illuminations (2 ms/s) and camera exposure times (100 ms/s for for both LED and laser) were synchronized by a dedicated program written in LabVIEW. Background images were captured by the same camera at a 1 frame/s rate to be used for offline correction of raw images.

### Experimental protocol and post-operative care

Multi-modal optical imaging of the mouse brain surface started with a baseline period of 10 min, followed by the bilateral occlusion of the CCAs (2-vessel occlusion; 2VO) for 45 minutes to induce incomplete global forebrain ischemia. The carotid arteries were then released to allow reperfusion, which was monitored for an additional 75 minutes. The wounds were sutured, and the animals were transferred to an incubator cage (32 °C) for 2 hours and allowed to wake up. Wounds were treated with Betadine (10 mg/ml, Egis) and Lidocaine (10 mg/ml, Egis) for local analgesia. Animals were hydrated by the administration of warmed (37 °C) saline (2 ml) subcutaneously. For long term analgesia, mice received a subcutaneous injection of the non-steroid anti-inflammatory drug Carprofen (5 mg/kg; Rycarfa, 8501 Novo Mesto, Slovenia). Finally, animals were supplied with fresh water and a softened standard chow in a recovery cage for 24 hours.

### Pharmacologic SD blockade

To attenuate SD evolution, the non-competitive NMDA receptor antagonist MK801 (Sigma-Aldrich, St. Louis, MO, USA) was administered in a separate cohort (0.3 mg/kg, i.p., n=29), 30 min prior to ischemia induction.

### Invasive mean arterial blood pressure (MABP) measurements

In additional mice (Control: n=5, MK801: n=7), MABP was measured invasively over the experimental protocol through a catheter inserted into the left femoral artery. The MABP signal was digitized in a dedicated hardware and software environment (Biopac MP150 and Acknowledge 4.2, Biopac Systems Inc., USA). Systemic blood gas analysis was performed 20 minutes before recording. Physiological variables (MABP, arterial partial pressure of O_2_, CO_2_ and pH) were in the physiological range (Table S1-S2).

### Characterization of post-stroke neurological deficit

Neurological score was taken a day before (baseline) and 24 hours after stroke (Fig. 1). Sensorimotor function was evaluated with the Composite Garcia Neuroscore (GN) scale^18^. The total score for the GN test ranges from 0 (severe impairment) to 21 (no deficit). All test domains were performed in the same order for each animal.

**Figure 1.**
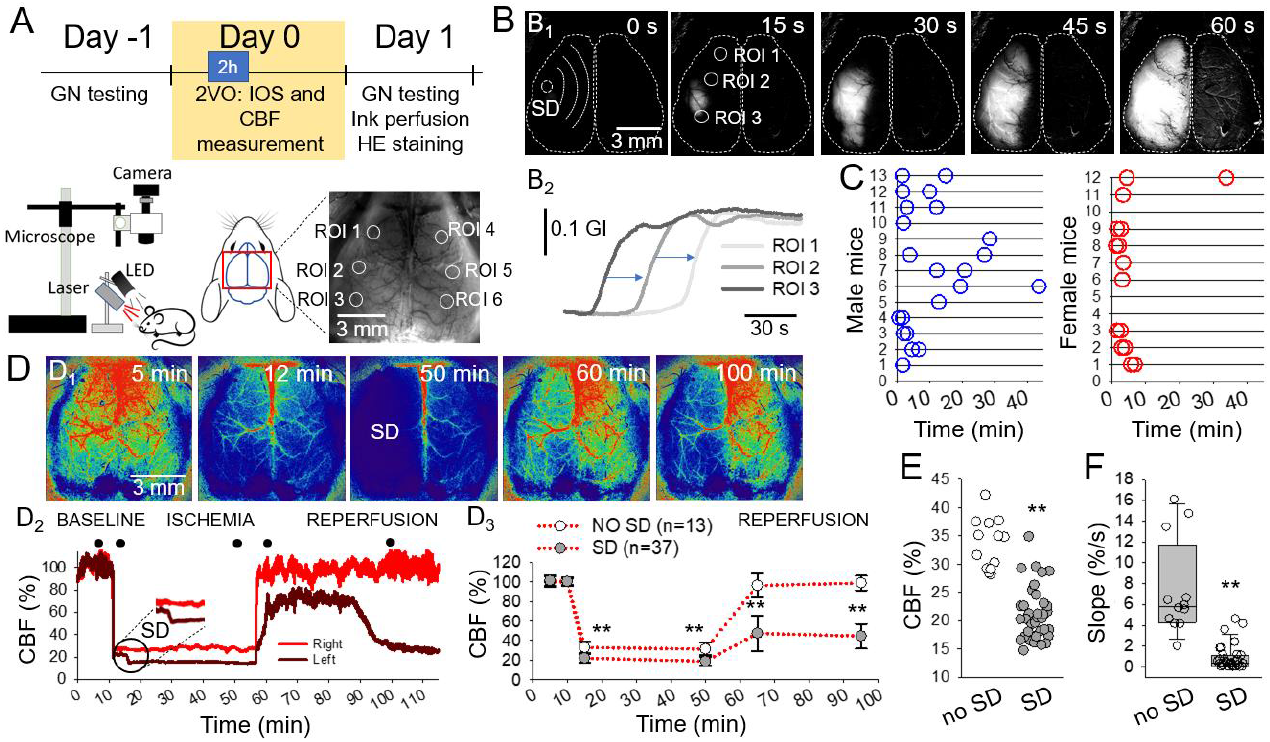
IOS and CBF imaging reveals spatial coincidence between the occurrence of SD during ischemia and the subsequent insufficiency of reperfusion.**A**, The timeline of the experiments, the arrangement of the imaging setup, and a representative raw transcranial IOS image of the mouse cortex with regions of interests placed offline (ROIs 1-6). **B**, A representative IOS image sequence of the cortical surface after background subtraction (B_1_) and corresponding IOS intensity traces (B_2_) demonstrate the unilateral occurrence of SD. **C**, The temporal pattern of first SD per hemisphere over the ischemic period (45 min) in male (n=13) and female (n=12) mice. **D**, Representative CBF maps (D_1_) of the same preparation as in B, and corresponding CBF traces (at ROI2 and ROI5 in A) (D_2_) show the spatial coincidence between SD occurrence and reperfusion failure. CBF images correspond in time with the closed circles over the CBF traces. Mean CBF values (D_3_) show significant reperfusion failure ipsilateral to ischemic SD at 15 and 45 min after the release of the carotid arteries. Note complete reperfusion in the hemisphere devoid of SD under ischemia. **E**, SD occurrence was linked to a greater drop of CBF after ischemia onset. **F**, Reperfusion ipsilateral to SD developed at a considerably slow rate (slope). Data are given as individual values (C & E), mean±stdev (D_3_) or in a box plot (F). The distribution of data was evaluated with a Shapiro-Wilk test (D_2_: p=0.065, E: p<0.050, F: p<0.050), followed by a One-Way RM ANOVA with Holm-Sidak post-hoc test (D_3_) and a Mann-Whitney Rank Sum Test (E&F), p<0.01**.

### Evaluation of the Circle of Willis (CoW) anatomy

Mice were transcardially perfused under deep anaesthesia with saline and 4% paraformaldehyde (PFA), followed by carbon black ink. The brains were carefully removed, and the CoW was evaluated with a stereomicroscope (Alpha STO-4M, Elektro-Optika Kft., Hungary) equipped with a Nikon-DS Fi3 camera.

### Hematoxylin-eosin staining to assess cell damage

After the anatomical evaluation of the CoW, 10 control and 10 MK801-treated brains were cut to 10-μm thick frozen sections with a freezing microtome (Leica CM 1860 UV, Leica, Germany). The sections were stained with hematoxylin and eosin (Sigma-Aldrich, USA) and cover slipped with Eukit® (Merck, USA). Photomicrographs of the sections were taken at 40× magnification with a Nikon-DS Fi3 camera attached to a Leica DM 2000 Led light microscope (Leica Microsystems GmbH, Germany).

### Data analysis and statistics

Power analysis for pharmacological experiments followed established principles. The pilot experiments indicated differences between the experimental groups (MK801 vs. Control) and achieved the confidence level of 95% and a power of 80% at low sample size (α = 0.05 and β (type II error) of 0.2). As calculated, sufficient statistical power was assumed at a final sample size of 11-12 animals/group. The calculations were run in GPower 3.1 (Heinrich Heine University of Düsseldorf, Germany).

All mice were coded independent of treatment or sex and randomly allocated to experimental groups. The four investigators performing surgeries, neuroscoring, reperfusion effficacy and infarct size calculation were completely blinded to treatment and animal gender. Six animals were excluded because ischemic injury was lethal during recording (n=2) or overnight to Day 1 (n=4).

Raw image sequences were analyzed in ImageJ (National Institute of Health, Bethesda, USA) and AcqKnowledge 4.2.0 (Biopac Systems Inc., USA). Local changes in IOS intensity and CBF with time were extracted by placing regions of interest (ROIs; 50 × 50 pixels, 0.4 × 0.4 mm) at selected sites in the images. CBF recordings obtained by LASCA were expressed relative to baseline by using the average CBF of the first 5 minutes of baseline (100%) and the recorded biological zero after terminating each experiment (0%). Neuronal necrosis was evaluated using the inbuilt automatized Plug-in “analyze particles” in ImageJ. Data are given as mean±stdev. Each data set was first evaluated with a Shapiro-Wilk test of normality to guide the choice of parametric or non-parametric statistics. Statistical analysis was conducted with the software SigmaPlot 12.5 (Systat Software, Inc., San Jose, CA, USA). Distinct statistical methods are provided in detail in each Figure legend.

## Results

### Spatial coincidence between SD during ischemia and subsequent reperfusion failure

Transcranial IOS imaging revealed the occurrence and propagation of SD over the mouse cerebral cortex during ischemia (Fig. 1A-B). Three distinct SD patterns were observed: (i) SD evolved in each hemisphere (bilateral SD, n=9/6, male/female; the two SDs emerged at unrelated foci in the two hemispheres and were apart in time), (ii) SDs predominantly occurred in one hemisphere (unilateral SD, n=4/3, male/female), or (iii) SD was not seen in either of the hemispheres (no SD, n=0/3, male/female). Altogether, 37 SDs were recorded in 25 mice. SD occurrence was slightly more likely in males than in females (hemispheres implicated in SD evolution: n=22/15, male/female).

CBF imaging indicated a sharp drop of CBF upon ischemia onset (Fig. 1D). A greater reduction of perfusion favored SD evolution (CBF drop to 21.1±4.6 vs. 33.6±4.4 %, SD vs. no SD) (Fig. 1E). In case SD occurred, a further flow reduction known as spreading ischemia was coupled to SD (to 18.6±4.3 vs. 31.2±6.3 %, SD vs. no SD) (Fig. 1D). Upon the release of the CCAs, reperfusion failure was seen in the hemisphere that had been engaged in SD (47.2±18.3 and 44.4±12.5 %, 15 min and 45 min after reperfusion initiation). In contrast, perfusion recovered to pre-ischemic levels in case the hemisphere had not undergone SD (96.3±11.9 and 98.9±7.4 %, 15 min and 45 min after reperfusion initiation) (Fig. 1D). Reperfusion failure after SD was also reflected by the low rate of flow elevation after the release of the carotid arteries, in contrast with the sharp rise of CBF in the hemisphere devoid of prior SD (0.87±1.19 vs. 7.45±4.60 %/s, SD vs. no SD) (Fig. 1F).

### The link between incomplete CoW and greater CBF reduction favoring SD

The CoW is often incomplete in young adult C57BL/6 mice^19^. The P1 segment of the posterior cerebral artery (PCA) is frequently absent or dysplastic^20^. Accordingly, the absence of the P1 segment in our mouse cohort was observed regularly (Fig. 2). Importantly, the absence of P1 predicted greater perfusion drop in the ipsilateral hemisphere (to 21.3±5.0 vs. 30.8±6.3 %, P1 absent vs. P1 present), which generally coincided with the occurrence of SD and subsequent reperfusion failure (Fig. 2D). The presence of P1 was seen more often in female mice (Fig. 2B).

**Figure 2.**
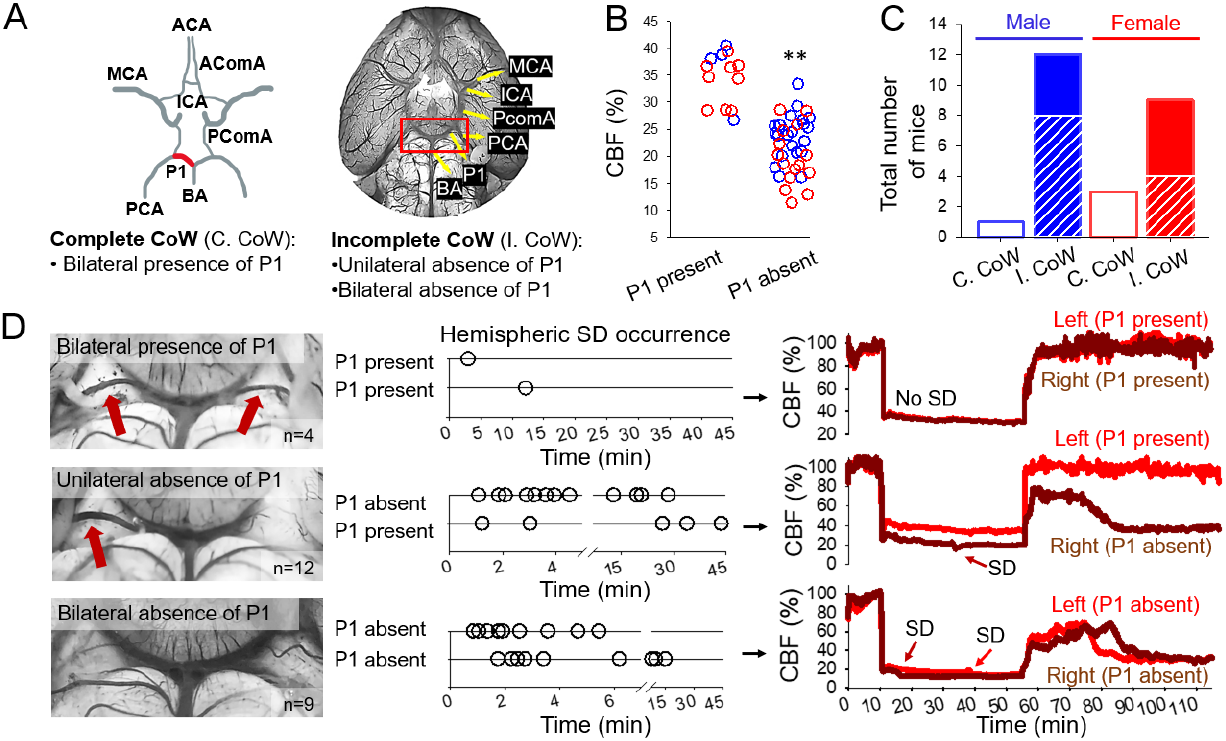
The absence of the P1 segment of the posterior cerebral artery (PCA) of the CoW predicts greater perfusion drop in response to CCA occlusion and subsequent SD occurrence.**A**, The anatomy of the CoW (P1 segment in red) represented schematically along an original photomicrograph of the base of a mouse brain, vasculature filled with carbon ink. **B**, CBF minimum measured shortly after ischemia induction in each hemisphere at the presence or absence of P1. Note that the absence of P1 is associated with lower CBF minimum (blue: male, red: female). **C**, The prevalence of P1 anatomical variations in the CoW in male and female mice. Complete CoW (C.CoW) is shown as open bars, while incomplete CoW (I.CoW) is represented by dashed portions for the unilateral and closed portions for the bilateral absence of P1. **D**, Selected photomicrographs demonstrate anatomical variations of the CoW. Red arrows show P1. Corresponding timelines indicate the temporal pattern of SDs detected in all our mice during 45 min ischemia. CBF traces depict representative reperfusion kinetics corresponding to the presence/absence of P1. Note that SDs under ischemia and reperfusion failure developed most frequently in the hemisphere ipsilateral to the absence of P1. Abbreviations: ACA, anterior cerebral artery; AComA, anterior communicating artery; BA, basilary artery; ICA, internal carotid artery; MCA, middle cerebral artery; PComA, posterior communicating artery. The distribution of data (panel B) was evaluated with a Shapiro-Wilk test (p=0.483), followed by a One-Way ANOVA, p<0.01**.

The bilateral presence of P1 was seen in 4 mice (n=1/3, male/female), in which only 2 SDs were detected. The unilateral absence of P1 in 12 mice (n=9/4, male/female) was invariably associated with the occurrence of SD in the ipsilateral hemisphere. In some of these mice (n=4/1, male/female), SD also occurred at the contralateral side in case the CBF in the contralateral cortex fell under the CBF threshold of SD initiation (Fig. S1). Finally, the bilateral absence of P1 in 9 mice (n=4/5, male/female) typically gave rise to SD in both hemispheres. The lower likelihood for SD to develop in female mice (Fig. 1C) appeared to be linked to the higher incidence of a complete CoW (CoW: 4/12 vs. 1/13, female vs. male) (Fig. 2C).

### Neurological dysfunction and neuronal injury associated with reperfusion failure

Mice that suffered SD and reperfusion failure displayed considerable somatosensory neurological deficit as expressed on the GN scale a day after ischemia (14.5±3.1 vs. 18.8±0.8, SD vs. no SD) (Fig. 3B). Forelimb outstretching, lateral turning, and climbing were linked to SD occurrence and reperfusion failure (Fig. 3B_1_). In contrast, mice that experienced no SD and had adequate reperfusion performed close to their pre-ischemic, optimal function (18.8±0.8 vs. 21.0, post-vs. pre-ischemia) (Fig. 3B). Further, higher composite GN score correlated closely with a higher level of reperfusion (Fig. 3C). Finally, neuronal necrosis was more prominent ipsilateral to SD and reperfusion failure, in all brain regions analyzed (e.g. parietal cortex: 130±2 vs. 97±13 cell count/1000 μm^2^, SD vs. no SD) (Fig. 3D).

**Figure 3.**
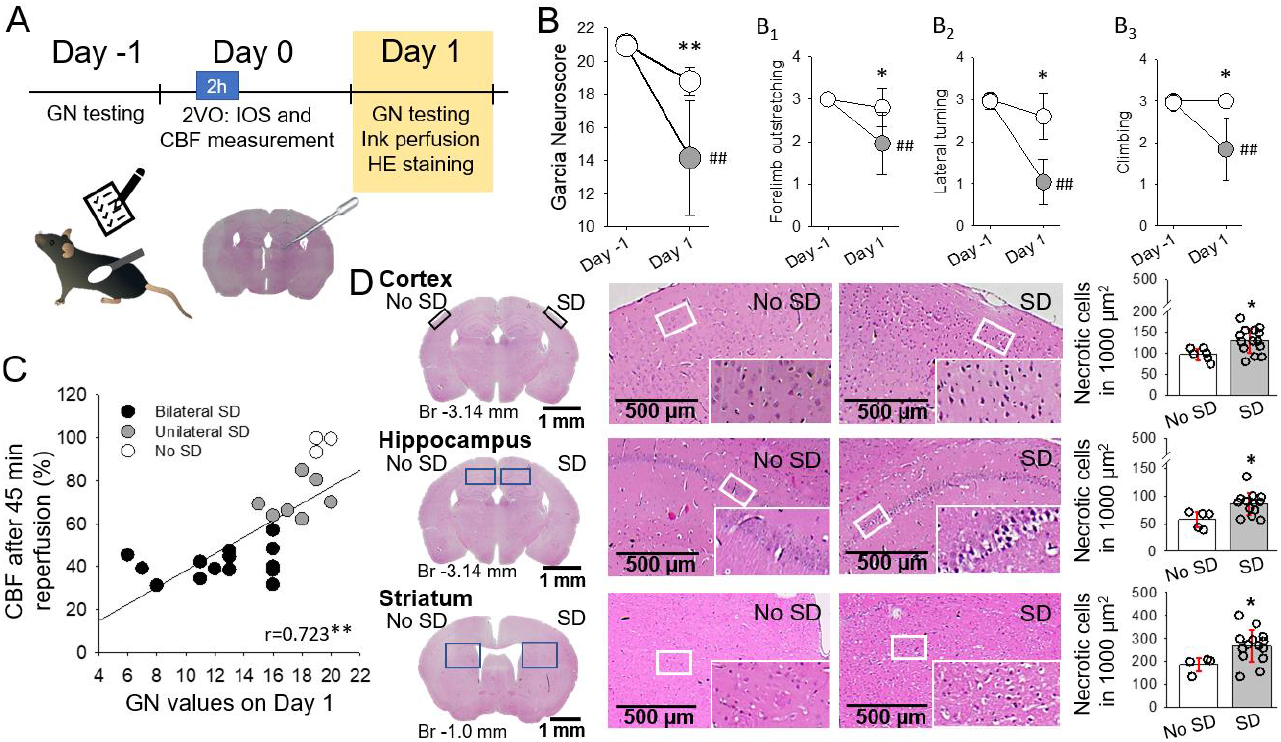
The occurrence of SD during ischemia and subsequent reperfusion failure are associated with severe neurological deficit and marked neuronal necrosis 24 h after ischemia.**A**, The timeline of the experiments. **B**, Neurological sensorimotor deficit expressed on the Garcia Neuroscore (GN) scale a day after ischemia, with respect to pre-ischemic performance. Selected motor test results revealed asymmetric motor function linked to SD occurrence. **C**, Positive linear correlation between GN score and reperfusion failure taken 45 min into reperfusion. **D**, Neuronal necrosis in hematoxylin-eosin (HE) stained sections. In B and D, data are given as mean±stdev. The distribution of data was evaluated by a Shapiro-Wilk test (B, p=0.599, B_1_, p<0.050. B_2_, p<0.050, B_3_, p<0.050, C, p=0.281, D, p=0.200), followed by a One-Way RM ANOVA with Holm-Sidak post-hoc test (B), a Kruskal-Wallis ANOVA (B_1_, B_2_, B_3_), a Two-Tailed Pearson correlation analysis (C) or one-way ANOVA (D), p<0.05* and p<0.01** vs. No SD, p<0.01^##^ vs. Day -1.

### Pharmacological SD inhibition to prevent reperfusion failure

AT and ÁM had full access to all the data in the study and takes responsibility for its integrity and the data analysis.

NMDA receptor antagonists inhibit but not completely block SD^21^. By the application of MK801, we set out to confirm that SD – rather than an incomplete CoW by itself – was implicated in reperfusion failure. As expected, MK801 curtailed SD, reflected by the smaller relative SD area (76.1±10.4 vs. 87.7±6.5 %, MK801 vs. vehicle) and the lower rate of SD propagation (3.2±1.3 vs. 4.2±1.4 mm/min, MK801 vs. vehicle) (Fig. 4A). The smaller SD area in the MK801-treated group offered the opportunity to test the efficacy of reperfusion at a cortical region unaffected by SD in addition to the area covered by SD within the same hemisphere (Fig. 4B_1_). As shown earlier, reperfusion was impaired in the cortical area affected by SD; yet reperfusion was optimal in the same hemisphere in the area which had not been invaded by SD (78.8±17.6 vs. 49.2±1.0 %, SD non-affected area vs. SD affected area) (Fig. 4B_1-2_). In line with improving reperfusion via SD inhibition, the MK801-treated mice achieved higher score on the functional test a day after ischemia (GN score: 17.5±1.9 vs. 14.6±2.5, MK801 vs. vehicle), although neuronal necrosis was not attenuated (Fig. S1).

**Figure 4.**
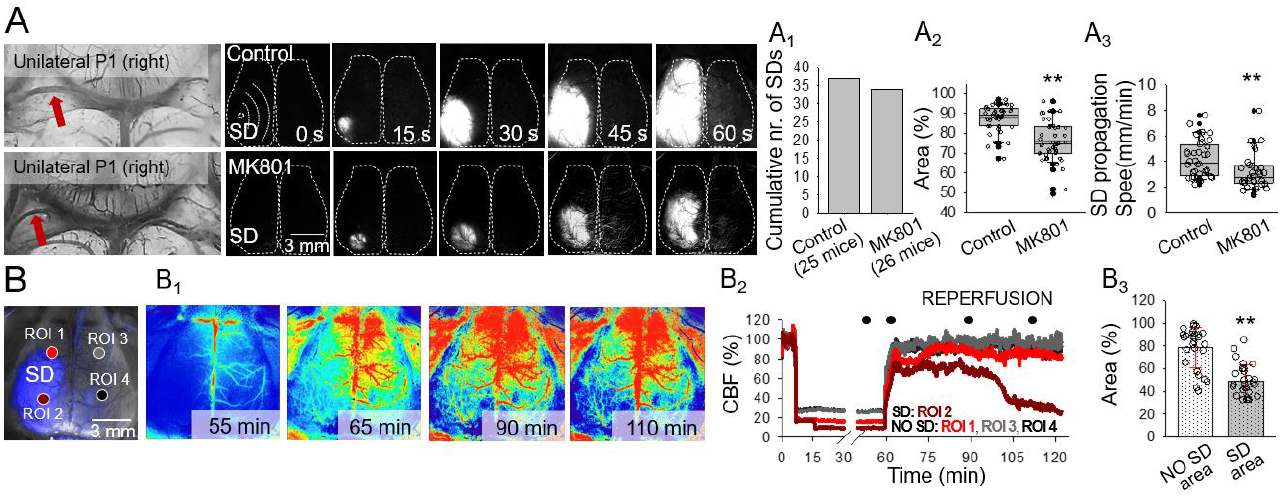
The pharmacological inhibition of SD counteracts reperfusion failure.**A**, Representative images demonstrate the absence of the left P1 at the base of the brain, and the corresponding evolution of SD in background subtracted IOS intensity images. Note that SD covered the entire ipsilateral cortex in a control mouse, while SD propagated a shorter distance in an MK801-treated preparation. MK801 did not block SD occurrence (A_1_) but reduced the area of the cortical surface bearing SD (A_2_) and decelerated SD propagation (A_3_). **B**, An IOS image demonstrates the cortical surface. In the left cortex, the area covered by SD is shown in blue. ROI1 was positioned over the surface unaffected by SD. In contrast, ROI2 was located within the area engaged in SD. ROI3-4 was placed over the contralateral cortex. The corresponding traces (B_1_) and the quantitative analysis (B_2_) reveal that reperfusion failure developed in the area previously engaged in SD (“SD area”), but reperfusion was sufficient in the same hemisphere in an area, which had not been reached by SD (“No SD area”). Data are given as mean±stdev. The distribution of data was evaluated by a Shapiro-Wilk test (A_2_, p=0.123, A_3_, p<0.050, B_3_, p=0.050), followed by a Mann-Whitney Rank Sum test (A_2_ and A_3_) or a T-test (D_3_), p<0.05*, p<0.01**.

## Discussion

This is the first study to demonstrate the decisive role of SD in reperfusion failure following cerebral ischemia. Ischemic SD evolution emerges as a reliable predictor of inadequate reperfusion despite successful restoration of large vessel patency. This is a significant realization, because in almost half of stroke patients treated with endovascular therapy or thrombectomy, recanalization is futile, and functional outcome is poor despite optimal angiographic results^2,3^.

Indeed, technically successful recanalization does not always ensure sufficient reperfusion or therapeutic benefit in ischemic stroke care^2,3^. The unexpected inefficacy of reperfusion therapy is attributed in part to no-reflow. No-reflow manifests at the level of the microvascular network. Despite full recanalization of a large vessel, microcirculatory flow is not restored due to the compromised patency of small arterioles and capillaries^22,23^. Interestingly, SD-related vasoconstriction has been recently shown to originate in the capillary bed in the mouse brain^24^. It is plausible, therefore, that SD imposes decisive, persistent microvascular constriction and possibly capillary stall events implicated in reperfusion failure.

The CBF response to SD is shaped by the actual balance between vasodilator and vasoconstrictor substances released in response to SD^25^. Under severe ischemia, vasoconstriction may override the response, which is known as spreading ischemia^26^, and has also been observed here (Fig. 1D_2_). The SD related vasoconstriction is governed by the high extracellular concentration of K^+27^ and accumulating arachidonic acid derivatives^28^ including 20-hydroxyeicosatetraenoic acid (20-HETE)^29^ and prostaglandin F2α^30,31^. Moreover, 20-HETE was shown to impair vasodilation evoked by neuronal activation in experimental focal cerebral ischemia^32^, which may be analogous with the depression of dilation to SD in ischemic brain tissue^33,34^. Taken together, we suggest that early reperfusion failure linked to SD here is possibly mediated by the accumulation of vasoconstrictive substances in the wake of SD, sustained during the early phase of reperfusion. We are currently exploring this possibility experimentally in detail.

Another significant observation of this study is that SD occurrence was associated with a critically low CBF related to reduced collaterals of the CoW. Anatomical anomalies of the CoW are common in the C57BL/6J mouse strain^19^, as well as in the healthy adult human population^35^. The absence or hypoplasia of the P1 segment of the PCA, the second most frequent vascular attenuation in human CoW anomalies^35^ caused in our mice a significantly greater drop of CBF after CCA occlusion. The drop of CBF below a critical level was followed by SD occurrence, which corresponds with the concept of a hypoperfusion threshold of SD generation^34^. In fact, low collateral grade in stroke patients correlates with larger final infarct volumes and worse long-term neurologic deficit^36^.

Both male and female mice were used in this study without expecting sexual dimorphism in the read-outs^37^. At close inspection of the data, we have found a lower incidence of SD in female mice. The lower SD frequency corresponded to a higher incidence of a complete CoW predicting a smaller CBF drop after vascular occlusion in female mice. Therefore, the better collateral circulation itself appeared to account for the fewer SDs – rather than any potential sex-related factor concerning ischemic tolerance or SD susceptibility.

In conclusion, we put forward the novel concept that SD is a main contributor to reperfusion failure despite successful recanalization after stroke. Moreover, our findings suggest that poor collaterals lead to a more severe drop of blood flow after cerebrovascular occlusion, which creates favorable conditions for SD initiation. We propose that the underlying mechanism of reperfusion failure due to SD is sustained vasoconstriction, which may manifest primarily at the level of the microvascular bed. Finally, the reperfusion failure after SD appears to predict aggravated neurological deficit and poor outcomes after stroke. Understanding the cause of futile recanalization and identifying SD as an underlying factor may guide new therapeutic options to improve stroke outcomes.

## Abbreviations

2VO: Bilateral common carotid artery occlusion (“two-vessel occlusion”)
20-HETE: 20-Hydroxyeicosatetraenoic acid
ACA: Anterior cerebral artery
AComA: Anterior communicating artery
BA: Basilary artery
C. CoW: Complete circle of Willis
CBF: Cerebral blood flow
CCA: Common carotid artery
CoW: Circle of Willis
GN: Garcia Neuroscore
HE: Hematoxylin-eosin
I. CoW: Incomplete circle of Willis
ICA: Internal carotid artery
IOS: Intrinsic optical signal
LASCA: Laser speckle contrast analysis
MCA: Middle cerebral artery
NMDA: N-methyl D-aspartate
PCA: Posterior cerebral artery
PComA: Posterior communicating artery
ROI: Region of interest
SD: Spreading depolarization

---

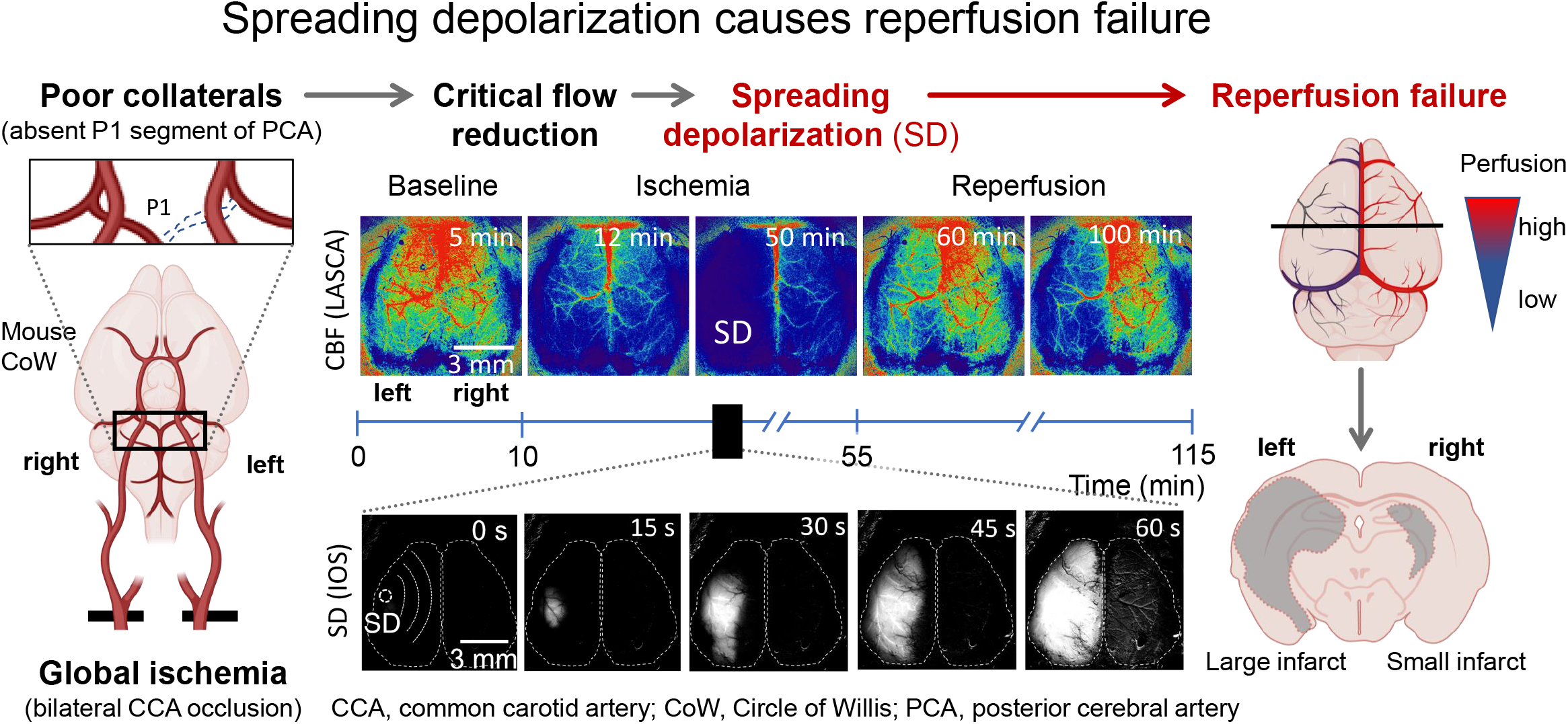

