## Supplementary material for "Spreading depolarization causes reperfusion failure after cerebral ischemia"

**Table S1.** Mean arterial blood pressure (MABP) changes over the experimental protocol.

| Group | n | Body weight (g) | MABP (mmHg) | | | | |
|---|---|---|---|---|---|---|---|
| | | | Baseline | 1 min after MK801 or vehicle injection | 1 min after 2VO induction | 1 min before reperfusion | 15 min after reperfusion |
| Control | 5 | 27±5 | 86±11 | 84±7 | 105±2 | 77±7 | 82±5 |
| MK801 | 7 | 26±5 | 85±7 | 85±5 | 104±14 | 79±9 | 81±7 |

Abbreviations: 2VO, bilateral common carotid artery occlusion, "two-vessel occlusion"

**Table S2.** Arterial blood gas analysis prior to ischemia induction.

| Group | n | Blood pH | $pO_2$ | $pCO_2$ |
|---|---|---|---|---|
| Control | 5 | 7.4±0.5 | 148±17 | 29±6 |
| MK801 | 7 | 7.4±0.3 | 132±13 | 28±9 |

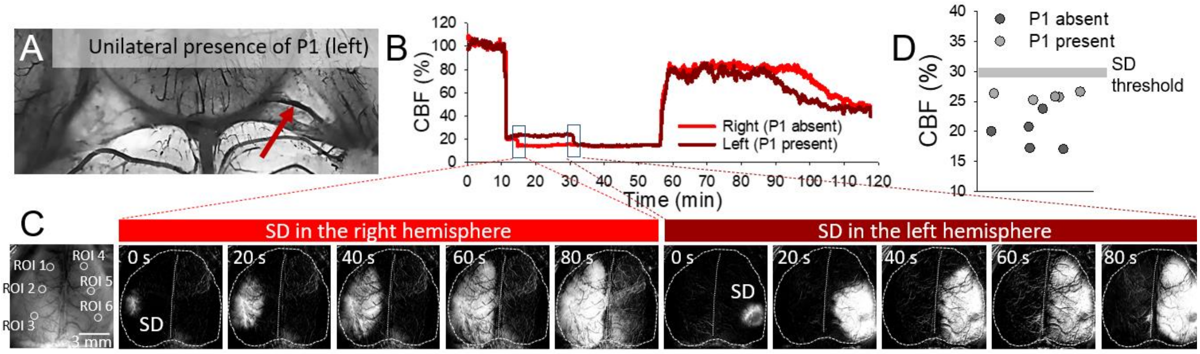


**Figure S1.** Irregular pattern of SD occurrence at the unilateral absence of P1.
**A,** A photograph of a circle of Willis, in which the P1 segment was absent on the right side and present on the left side (red arrow). **B,** CBF traces from the two hemispheres corresponding to panel A. Note that even though the P1 was present on the left side, the drop of CBF after bilateral CCA occlusion was equally deep in both hemispheres, diving below the CBF threshold of SD. **C,** SD occurred first in the right hemisphere (P1 absent), and later in the left hemisphere (P1 present), as shown in background subtracted IOS intensity images. **D,** In 5 mice (n=4/1, male/female), SD occurred first in the hemisphere ipsilateral to the absence of P1, and then in the contralateral hemisphere, where P1 was present. In all these mice, CBF dropped in both hemispheres under the CBF threshold of SD.

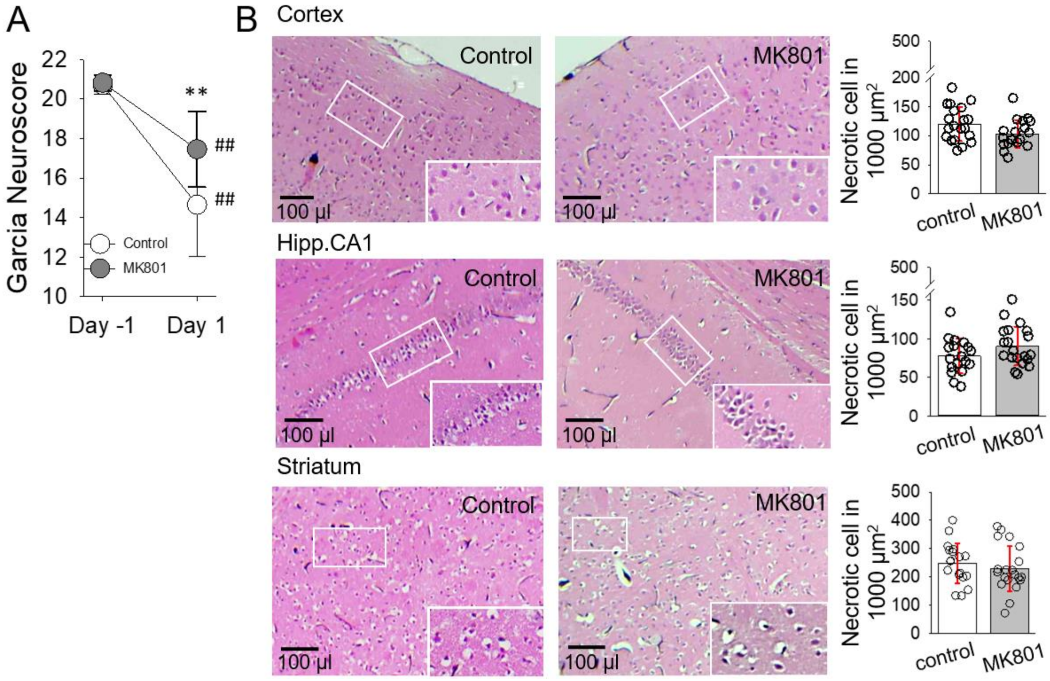


**Figure S2.** The effect og MK801 treatment on neurologic function and necrotic cell deat 24 h after ischemia. **A,** Garcia Neuroscore values a day before (Day -1) and a Day after (Day 1) with respect to ischemia. **B,** Hematoxylin-eosin stained representative coronal brains slices and quantitative evaluation show that MK801 treatment was ineffective on necrotic cell death. Data are given as mean±stdev. The distribution of data was evaluated by a Shapiro-Wilk normality test (A, p=0.876; B, p<0.05) followed by an RM ANOVA with a Holm-Sidak posthoc medhod (A; p<0.01** vs. Control, p<0.01$^{##}$ vs. Day -1) or a ranks Dunn's posthoc method (B; no significant differences: cortex, p=0.980; CA1, p=0.108; striatum, 0.774.
